# Molecular characterization of a novel cytorhabdovirus with a unique genomic organization infecting yerba mate (*Ilex paraguariensis*) in Argentina

**DOI:** 10.1101/2020.01.28.923201

**Authors:** Nicolás Bejerman, Raúl Maximiliano Acevedo, Soledad de Breuil, Oscar A. Ruiz, Pedro Sansberro, Ralf G. Dietzgen, Claudia Nome, Humberto Debat

## Abstract

The genome of a novel rhabdovirus was detected in yerba mate (*Ilex paraguariensis* St. Hil.). The newly identified virus, tentatively named yerba mate virus A (YmVA), has a genome of 14,961 nucleotides. Notably, eight open reading frames were identified in the antigenomic orientation of the negative-sense, single-stranded viral RNA, including two novel accessory genes, in the order 3’-N-P-3-4-M-G-L-8-5’. Sequence identity of the encoded proteins as well as phylogenetic analysis suggest that YmVA is a new member of the genus *Cytorhabdovirus*, family *Rhabdoviridae*. YmVA unique genomic organization and phylogenetic relationships indicate that this virus likely represents a distinct evolutionary lineage within the cytorhabdoviruses.

---

Yerba mate (*Ilex paraguariensis* St. Hil., Aquifoleaceae) is a subtropical tree cultivated in the northeast of Argentina, south of Brazil, and the west of Paraguay. Its leaves and stems are widely utilized in the preparation of an infusion popularly known as “mate.” In Argentina, the main producer, the cultivated area has reached 165,327 ha [1]. Although yerba mate is of significant economic, social and traditional importance in the region, studies on plant diseases are still scarce, and only a handful of viruses have been identified in this holly species [2-4]

Recently, the complete genome sequence of another cytorhabdovirus was described, associated with chlorotic linear patterns, chlorotic rings, and veins yellowing symptoms in yerba mate, which was named yerba mate chlorosis-associated virus (YmCaV) [3]. The genus *Cytorhabdovirus* in the family *Rhabdoviridae* has the largest number of members among the four genera of plant-infecting viruses in the family. Cytorhabdoviruses have a unsegmented, negative-sense, single-stranded RNA genomes with six to ten open reading frames (ORFs) with a canonical genomic organization that includes six proteins in the order 3′-nucleocapsid protein (N) - phosphoprotein (P) - (putative) cell-to-cell movement protein - matrix protein (M) –glycoprotein (G) – polymerase (L) −5′ [5]. In this work, we report the molecular characterization of a novel and distinct cytorhabdovirus associated with yerba mate in Argentina, which we tentatively named yerba mate virus A (YmVA).

The raw sequence data analyzed in this study correspond to an RNAseq library (SRA: SRP110129), associated with the National Center for Biotechnology Information (NCBI) Bioproject PRJNA375923. Data were obtained by Illumina Hiseq 1500 sequencing of total RNA isolated from yerba mate mature leaves collected in Gobernador Virasoro, Corrientes, Argentina (BioSample: SAMN07206716), that did not show any visible virus-like symptoms [6]. The 390,909,798 2×101nt raw reads from the SRA were computationally analyzed according to Debat and Bejerman [7]. In brief, the raw reads were trimmed and filtered using the Trimmomatic tool v 0.39 (http://www.usadellab.org/cms/index.php?page=trimmomatic) and *de novo* assembled using Trinity v2.8.6 (https://github.com/trinityrnaseq/trinityrnaseq/releases). The resulting contigs were subjected to BLASTX searches (E-value < 1e^-5^) against the complete Refseq release of virus proteins (ftp://ftp.ncbi.nlm.nih.gov/refseq/release/viral/). A rhabdovirus-like 14,945 nt long contig was detected showing only 44.9% overall sequence identity to YmCaV. This contig was supported by a total of 21,012 reads (mean coverage = 140.5X). To confirm the obtained sequence, reverse transcription PCR (RT-PCR), cloning, and Sanger sequencing using specific primers designed from the assembled sequence were used; genome termini were obtained using the 3′ and 5′ RACE system for Rapid Amplification of cDNA ends (Life Technologies). Sequences were assembled and analyzed using Geneious v.8.1.9 (Biomatters Ltd).

The complete genome of YmVA is 14,961 nt (GenBank accession MN781667) and contains eight ORFs that were predicted using ORFfinder (https://www.ncbi.nlm.nih.gov/orffinder/, minimal ORF length = 150 nt) in the anti-genomic strand (Fig. 1). The molecular weight and isoelectric point of each YmVA encoded protein was calculated using the Compute pI/Mw tool available at ExPASy (https://web.expasy.org/compute_pi/) (Table S1). BlastP searches of the deduced proteins encoded by each ORF identified ORFs 1 and 7, as coding for the nucleocapsid protein (N) and RNA-dependent RNA polymerase (L), respectively, that are 30% and 46% identical with the aligned portion of partial N and L proteins encoded by the tentative cytorhabdovirus Iranian citrus ringspot-associated virus (IrCRSaV, [8]) (GenBank KU660038 and KU660039, respectively). None of the other predicted proteins had significant matches with any GenBank entries (Table S1). This may reflect the unavailability of the corresponding protein sequences of the closest hit, IrCRSaV, or any other rhabdovirus with the distinctive genomic organization of YmVA. The unique genome structure of YmVA shows coding sequences are flanked by 3’ leader (l) and 5’ trailer (t) sequences, that are 470 and 348 nt long, respectively, and a genome organization of 3’–N–P–3–4–M–G–L–8–5’ (Fig. 1). Interestingly, the YmVA genome and 3′ leader sequence are the longest described so far among cyto- and nucleorhabdoviruses (Table S2). The YmVA genome organization is different from all other cyto- and nucleorhabdoviruses [9], because of two novel accessory genes, tentatively named gene 3 and gene 8, based on their position in the genome. Accessory genes have been reported for several other animal and plant rhabdoviruses, located in various positions in the genome [9-10]. Nevertheless, to the best of our knowledge, YmVA is the first rhabdovirus identified so far which contains an accessory gene between L gene and 5′trailer [5-11]. YmVA is also the first cytorhabdovirus that has an accessory gene located between the P and putative movement protein genes. A gene in this location has been reported for the nucleorhabdovirus maize fine streak virus (MFSV) [12] (Table S2). The predicted YmVA protein 3 has 61 amino acids (aa) with a basic isoelectric point (IEP) of 7.96 (Table S1), whereas MFSV protein 3 has 93 aa with an acidic IEP of 5.45. YmVA gene 8 encodes a protein of 141 aa with a predicted molecular weight of 16.63 kDa and an isoelectric point of 6.15 (Table S1). No known functional domains were identified in either YmVA proteins 3 or 8. Future studies will explore the potential functions of these unique putative gene products. Unexpectedly, YmVA ORF6, which likely encodes the viral glycoprotein (G) based on its location in the genome and the two transmembrane (TM) domains identified in its N-terminal and central region (aa positions 21-43, 48-70) (Table S1), is remarkably short (155 aa) and appears to be the smallest reported rhabdovirus glycoprotein (Table S2). Similar TM domains, predicted using TMHMM version 2.0 (http://www.cbs.dtu.dk/services/TMHMM/), were also reported in the G proteins of other plant rhabdoviruses [4, 13], consistent with their membrane-associated functions. A signal peptide was also predicted using SignalP 3.0 server (http://www.cbs.dtu.dk/services/SignalP-3.0/) in the ORF6 encoded protein. Similar signal peptides have been predicted in the glycoprotein sequences of several cytorhabdoviruses [14-16] lending supports to the notion that ORF6 may encode a glycoprotein. Furthermore, and again highlighting the distinctiveness of YmVA, its predicted P and L proteins are the longest so far among cyto- and nucleorhabdoviruses (Table S2).

**Figure 1.**
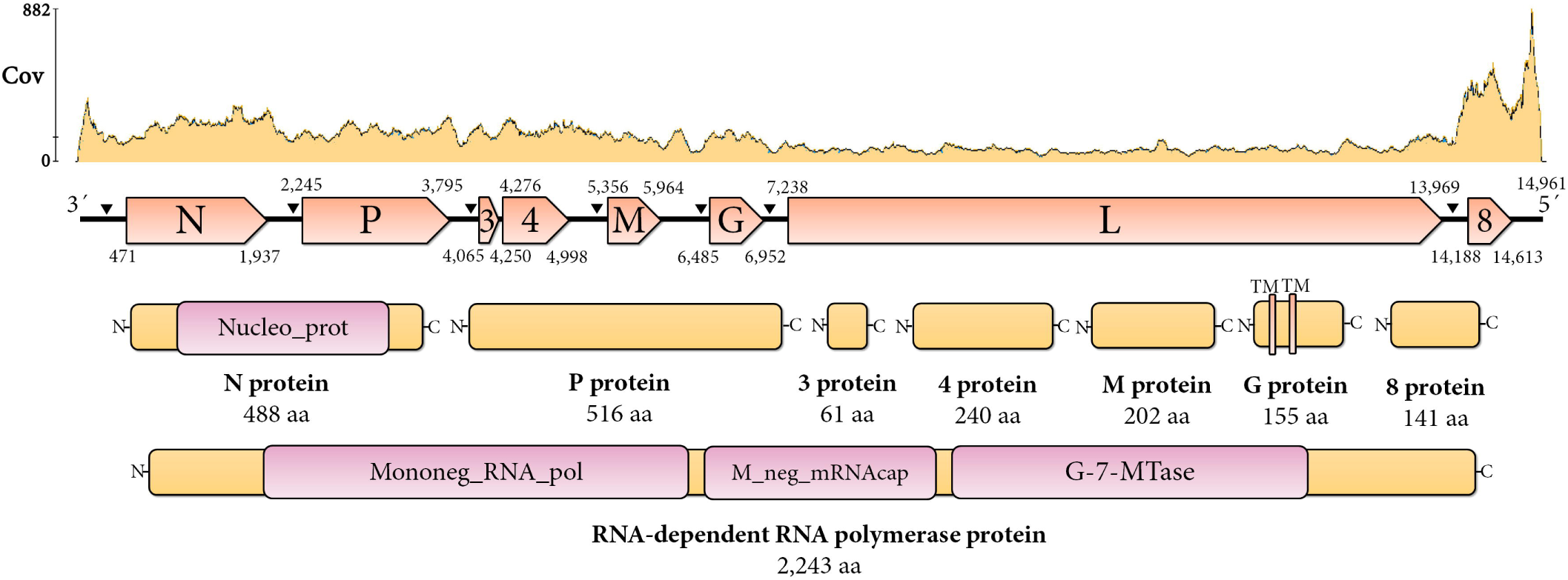
Genome graphs depicting coverage landscape, as number of reads supporting each position, architecture, and predicted gene products of YmVA. The predicted coding sequences are shown in orange arrow rectangles, start and end coordinates are indicated. Gene products are depicted in curved yellow rectangles and size in aa is indicated below. Predicted domains are shown in curved pink rectangles. Abbreviations: Cov, Coverage; N, nucleoprotein CDS; P, phosphoprotein CDS; 3, accessory protein 3 CDS; 4, putative cell-to-cell movement protein CDS; M, matrix protein CDS; G, glycoprotein CDS; L, RNA dependent RNA polymerase CDS; 8, accessory protein 8 CDS; TM, trans-membrane domain. Black triangles indicate locations of gene junctions.

Like all plant rhabdoviruses, YmVA ORFs are separated by intergenic “gene junctions” regions, which are composed of the polyadenylation signal of the preceding gene, a short intergenic non-transcribed region, and the transcriptional start of the following gene (Table S3). YmVA consensus “gene junction” region sequence is 3′ AUUCUUUUUGGUCCU 5′ and almost identical to that of the cytorhabdoviruses colocasia bobone disease associated virus (CBDaV) and papaya virus E (PpVE) (Table S3). This sequence is conserved across most gene junctions of YmVA, with only minor variations observed in the G-L and L-8 intergenic regions (Table S3). Unexpectedly, there was no identifiable, conserved “gene junction” sequence between gene 3 and gene 4 of the YmVA genome, like in the case of the other yerba mate cytorhabdovirus YmCaV [3], suggesting that these genes may be transcribed as a single transcript.

Amino acid sequence comparisons between the deduced YmVA proteins and the corresponding proteins of other cytorhabdoviruses revealed a very low sequence identity (data not shown). The highest aa sequence identity of less than 50% was with IrCRSaV, far below the species demarcation criteria for cytorhabdoviruses of 75% [5]. Low sequence identity is quite common between cytorhabdoviruses, that display a high level of diversity in both their genome sequence and organization [9]. Regarding all the other predicted gene products no significant hits were retrieved using BLASTP or PSI-BLAST. However, it is worth mentioning that using HHBlits with standard parameters (https://toolkit.tuebingen.mpg.de/tools/hhblits) we found a low similarity of YmVA protein 4 with the 4b protein of wuhan insect virus 6 (WhIV6) (E-value not significant). We mention this hit given that WhIV6 has been classified as a cytorhabdovirus and there is an apparent synteny in terms of genomic architecture for these two proteins. In addition, we found a hit of YmVA G protein with the systemic acute respiratory syndrome virus (*Coronaviridae*) envelope protein E (E-value not significant). We comment on this hit given the functional implications of this affinity. These tentative results should be taken as speculative and by no means indicate a confirmed link between the aforementioned proteins.

The genome sequence of YmVA, which has a unique organization of predicted ORFs among plant rhabdoviruses, illustrates the complexity of genome evolution of these viruses, where new accessory ORFs in previously undescribed positions are being identified. Supporting this view, a new cytorhabdovirus, named strawberry-associated virus 1 (SaV1), with a distinctive genome organization among rhabdoviruses, encoding not only one, but two accessory ORFs between G and L genes was recently identified in strawberry [17]. Furthermore, two nucleorhabdoviruses recently identified in alfalfa and apple, alfalfa-associated nucleorhabdovirus (AaNV) and apple rootstock virus A (ApRVA) also have a divergent genomic architecture among plant rhabdoviruses with an accessory ORF (U) located between M and G genes [18-19]. Thus, we suggest that future characterization of complete genome sequences of novel plant rhabdoviruses may result in the identification of additional novel accessory ORFs in unique positions between or within other genes, leading to a better understanding of rhabdovirus evolution.

To provide insights into the evolutionary history of YmVA, phylogenetic analysis was done, using all publicly available complete plant rhabdovirus polymerase sequences and the partial IrCRSaV L protein using MAFFT v7 (https://mafft.cbrc.jp/alignment/software/) with multiple aa sequence alignments using E-INS-i as the best-fit model. The aligned protein sequences were subsequently used as input in FastTree 2.1.11 (http://www.microbesonline.org/fasttree/) to generate phylogenetic trees by using the maximum-likelihood method (best-fit model = JTT-Jones-Taylor-Thorton with single rate of evolution for each site = CAT). Local support values were computed using the Shimodaira-Hasegawa test (SH) with 1,000 replicates. In the resulting phylogenetic tree, YmVA clustered in the genus *Cytorhabdovirus*, close to IrCRSaV apparently forming a distinct evolutionary lineage (Fig. 2). Among cyto- and nucleorhabdoviruses there is a strong correlation between phylogenetic relationships and the insect vector [9], so it is tempting to speculate that YmVA and IrCRSaV may be transmitted by a so far undescribed vector. Future studies should explore the potential functions of the novel rhabdovirus genes described here for YmVA, including comparisons with IrCRSaV the genome sequence of which is currently incomplete, and potential additional viruses from this intriguing lineage.

**Figure 2.**
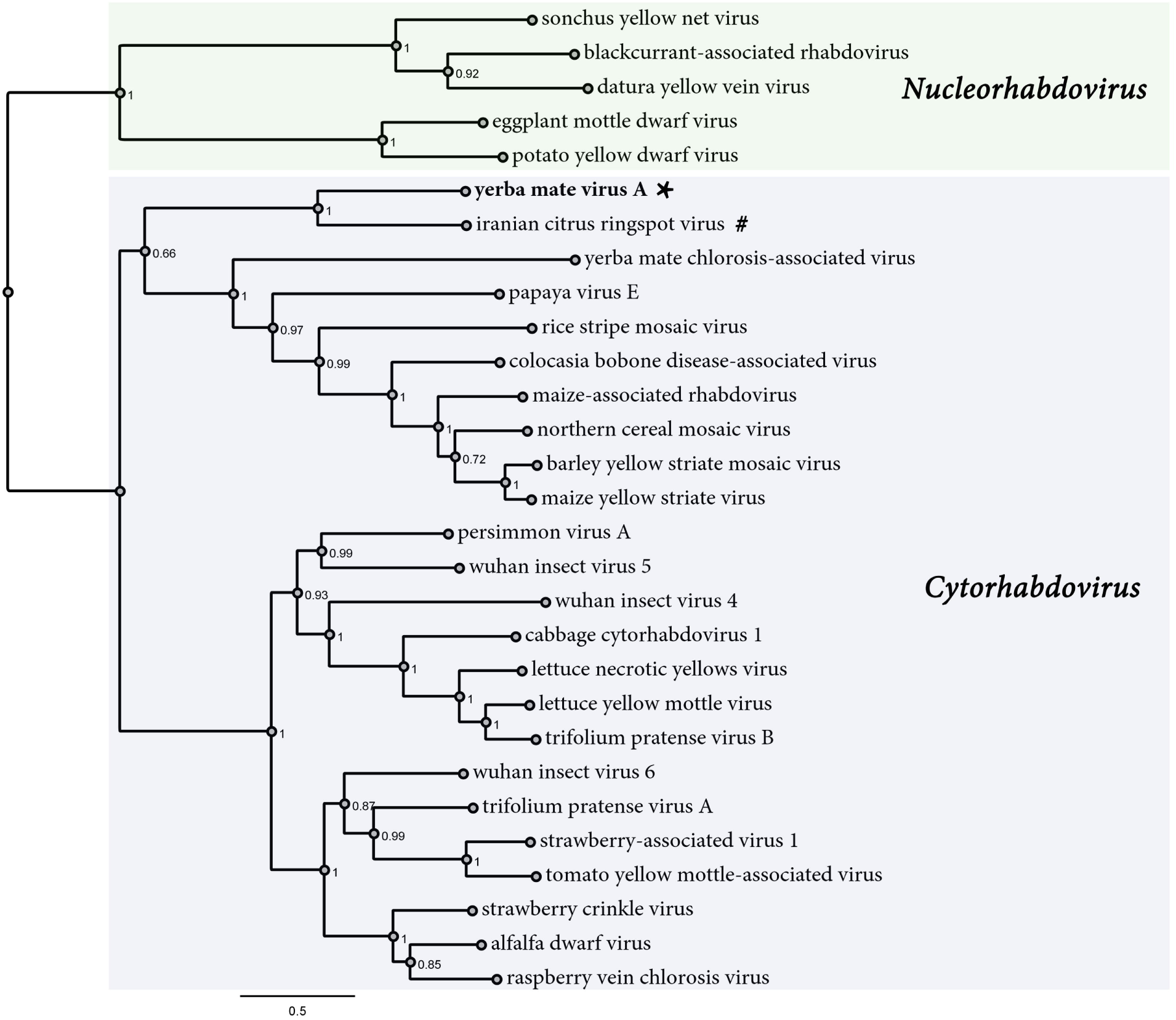
Maximum likelihood phylogenetic tree based on amino acid sequence alignments of the L polymerase of YmVA and other plant rhabdoviruses. The tree is rooted at the midpoint; nucleorhabdovirus and cytorhabdovirus clades are indicated by green and blue rectangles, respectively. The scale bar indicates the number of substitutions per site. Node labels indicate FastTree support values. The viruses used to construct the tree, and their accession numbers are: alfalfa dwarf virus (ADV; KP205452), barley yellow striate mosaic virus (BYSMV; KM213865), blackcurrant-associated rhabdovirus (BCaRV; MF543022), cabagge cytorhabdovirus 1 (CCyV1; KY810772), colocasia bobone disease associated-virus (CBDaV; KT381973), datura yellow vein virus (DYVV; KM823531), eggplant mottled dwarf virus (EDMV; NC_025389), lettuce yellow mottle virus (LYMoV; EF687738), lettuce necrotic yellows virus (LNYV; NC_007642), maize-associated rhabdovirus (MaCyV; KY965147), maize yellow striate virus (MYSV; KY884303), northern cereal mosaic virus (NCMV; AB030277), papaya virus E (PpVE; MH282832), persimmon virus A (PeVA; NC_018381), potato yellow dwarf virus (PYDV; GU734660), raspberry vein chlorosis virus (RVCV; MK257717), rice stripe mosaic virus (RSMV; MH720464), strawberry-associated virus 1 (SaV1; MK159261), strawberry crinkle virus (SCV; MH129615), sonchus yellow net virus (SYNV; L32603), trifolium pratense virus A (TpVA; MH982250), trifolium pratense virus B (TpVB; MH982249), tomato yellow mottle-associated virus (TYMaV; KY075646), wuhan insect virus 4 (WhIV4; KM817650), wuhan insect virus 5 (WhIV5; KM817651), wuhan insect virus 6 (WhIV6; KM817652), yerba mate chlorosis-associated virus (YmCaV; KY366322), iranian citrus ringspot-associated virus (IrCRSaV, KU660039). * indicates YmVA. # the IrCRSaV L protein used for this tree is truncated.

In conclusion, the unique genome organization, sequence identities and phylogenetic relationship indicate that YmVA should be considered as a representative of a new species in the genus *Cytorhabdovirus*, family *Rhabdoviridae*.

## Supporting information

Virus sequence MN781667

## Acknowledgments

This work was supported by ANPCyT (PICT 2014-1246), (PICT 2014-1212), (StartUp-PICT 2014-3648).

## Compliance with ethical standards

There is no conflict of interest

