## Supplementary Table 1 for "Molecular characterization of a novel cytorhabdovirus with a unique genomic organization infecting yerba mate (*Ilex paraguariensis*) in Argentina"

**Supplementary Table 1**. Characteristics of deduced proteins encoded by yerba mate virus A (YmVA) genome determined by predictive algorithms

| ORF No^a^ | Gene name | Calculated *Mr* (kDa) | isolelectric point | Putative Function | TMHMM | Highest scoring virus protein/*E*-value/query coverage (Blast P) |
| --- | --- | --- | --- | --- | --- | --- |
| 1 | N | 55.90 | 6.1 | nucleocapsid protein | none | IrCRSaV-N/  3e^-40^/30% |
| 2 | P | 59.10 | 5.82 | phosphoprotein | none | no hits |
| 3 | 3 | 6.84 | 7.96 | unknown | none | no hits |
| 4 | 4 | 26.8 | 8.63 | movement protein | none | WhIV6-4b/3.1/46%^b^ |
| 5 | M | 22.50 | 6.43 | matrix protein | none | no hits |
| 6 | G | 18.20 | 8.75 | glycoprotein | aa  21-43, 48-70 | SARS-E/180/36%^c^ |
| 7 | L | 259.20 | 6.79 | polymerase | none | IrCRSaV-L/0.0/46% |
| 8 | 8 | 16.6 | 6.15 | unknown | none | no hits |

^a^ ORF numbers are represented from 3´ to 5´ for genomic sense and correspond to those in Fig.1. ^b^ The hit of gene 4 ORF with the 4b protein of wuhan insect virus 6 (WhIV6) was obtained using HHBlits and its E-value is not significant. We mention this hit given that WhIV6 appears to be a cytorhabdovirus and the synteny in terms of genomic architecture of WhIV6 4b and YmVA ORF4. ^c^ The hit of G protein with severe acute respiratory syndrome (SARS) virus envelope protein E was obtained using HHBlits and its E-value is not significant. We show this hit given the functional implications of this affinity. These results should be considered as speculative. Iranian citrus ringspot-associated virus, IrCRSaV.

TMHMM=Transmembrane domain.
