## Supplementary Table 3 for "Molecular characterization of a novel cytorhabdovirus with a unique genomic organization infecting yerba mate (*Ilex paraguariensis*) in Argentina"

**Supplementary Table 3**. Gene junction regions of yerba mate virus A (YmVA) and other selected cytorhabdoviruses.

| YmVA | 3´end mRNA | intergenic sequence | 5´end mRNA |
| --- | --- | --- | --- |
| 3´leader/N | AUUCUUUUU | GGU | CCU |
| N/P | AUUCUUUUU | GGU | CCU |
| P/3 | AUUCUUUUU | GGU | CCU |
| 4/M | AUUCUUUUU | GGU | CCU |
| M/G | AUUCUUUUU | GGU | CCU |
| G/L | AUUUAUUUU | GGAU | CCU |
| L/8 | AUUAUUUUA | GGC | UCU |
| consensus | AUUCUUUUU | GGU | CCU |
| CBDaV | AUUCUUUUU | GG (N)n | CUC |
| PpVE | AUUCUUUUU | GGC | CCU |

YmVA sequences are highlighted in grey. For virus acronyms and GenBank accession numbers see legend of Fig.2. Note: There is no identifiable gene junction region between genes 3 and 4.
